# mmgenome: a toolbox for reproducible genome extraction from metagenomes

**DOI:** 10.1101/059121

**Authors:** Søeren M. Karst, Rasmus H. Kirkegaard, Mads Albertsen

## Abstract

**Summary:** Recovery of population genomes is becoming a standard analysis in metagenomics and a multitude of different approaches exists. However, the workflows are complex, requiring data generation, binning, validation and finishing to generate high quality population genome bins. In addition, several different approaches are often used on the same dataset as the optimal strategy to extract a specific population genome varies. Here we introduce mmgenome: a toolbox for reproducible genome extraction from metagenomes. At the core of mmgenome is an R package that facilitates effortless integration of different binning strategies by collecting information on scaffolds. Genome binning is facilitated through integrated tools that support effortless visualizations, validation and calculation of key statistics. Full reproducibility and transparency is obtained through Rmarkdown, whereby every step can be recreated.

**Availability and implementation:** The binning framework of mmgenome is implemented in R. Wrapper scripts for data generation and finishing is written in Perl. The mmgenome toolbox and associated step-by-step guides are available at http://madsal-bertsen.github.io/mmgenome/.

**Supplementary information:** Supplementary data are available at *Bioinformatics* online.

## 1 INTRODUCTION

Metagenomics, the sequencing of bulk DNA from a given microbial sample, is increasingly being used to recover population genomes directly from the environment. The process of separating metagenome contigs into population genomes is termed binning and rely on associating contigs with similar characteristics (Kunin *et al.*, 2008). For each contig, a number of different characteristics can be obtained: length, nucleotide frequency (e.g. GC or tetranucleotide frequency), coverage (metagenome abundance), paired-end read linkage, proximity information, presence of key genes and similarity to sequenced reference genomes.

In the past ten years a multitude of unsupervised metagenome binning methods have been developed, which often rely on a single characteristic analysed in different statistical frameworks (summarized by Mande *et al.*, 2012). Recently, approaches have been developed that leverage the dramatic decrease in sequencing cost, by taking advantage of differential coverage across a number samples (Nielsen *et al.*, 2014; Alneberg *et al.*, 2014). However, the unsupervised methods suffers from the fact that different strategies are often required for optimal binning of each genome, and subsequent manual refinement is needed to obtain high quality population genome bins.

The first metagenome binning approaches used supervised methods that integrated most of the raw contig characteristics to enable recovery of population genomes from simple communities (Tyson *et al.*, 2004). In addition, several supervised workflows have been described that integrate a number of different unsupervised methods at various stages of the binning process (Wrighton *et al.*, 2012; Sharon *et al.*, 2013; Albertsen *et al.*, 2013). However, supervised methods are often laborious and difficult to reproduce.

Here we present mmgenome: a toolbox for reproducible genome recovery from metagenomes. It provides a framework for integration of unsupervised methods in a transparent and reproducible supervised binning process. Thereby leveraging the shortcomings of both supervised and unsupervised methods.

## 2 DESCRIPTION

### 2.1 Data generation

The basic data needed by mmgenome is a metagenome assembly in fasta format and at least one associated coverage profile. If paired-end reads are available, the script network.pl can be used to generate a contig linkage file from a SAM file of the read mapping. A simple shell script is provided that integrates prodigal (Hyatt *et al.*, 2010), HMMER3 (Finn *et al.*, 2011), BLAST (Altschul *et al.*, 1990) and MEGAN (Huson *et al.*, 2011) to predict and taxonomic classify essential single copy genes, which can be used for both validation and visualization. Any additional information can be integrated as flat text files, e.g. taxonomic classification using Phy-loPythiaS+ (Gregor *et al.*, 2014) and 16S rRNA information.

**Fig. 1.**
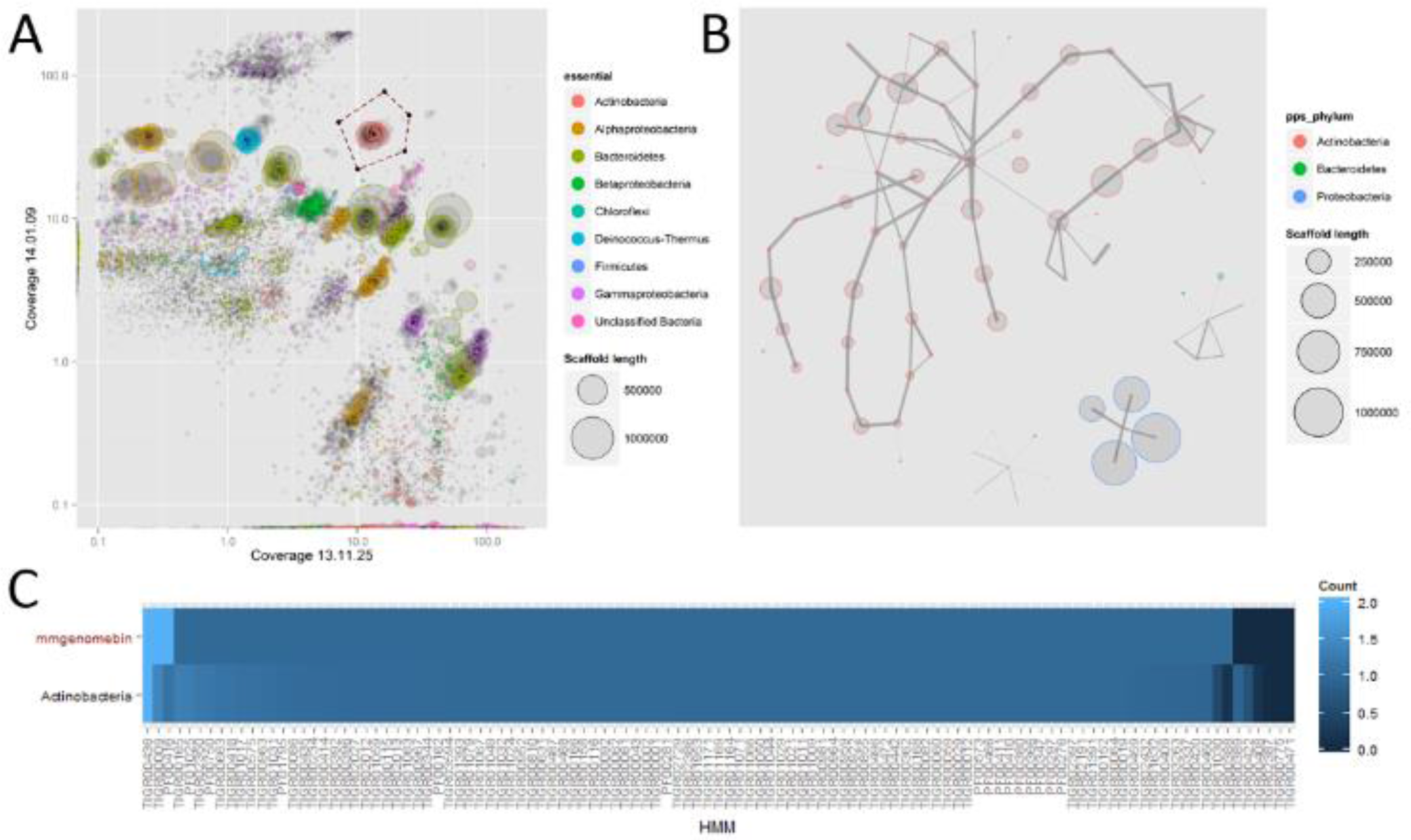
Example of individual genome extraction using the mmgenome R package. A) Differential coverage plot (*mmplot*) using of two different time points. Each scaffold is a circle, scaled by length and colored according to taxonomic classification of essential genes. A subset of the metagenome contigs is extracted interactively on the plot. B) A network plot *(mmplotnetwork)* of paired-end read connections between the extracted contigs is used to include repeats and identify contamination. C) Comparison of essential “single copy” genes with all sequenced Actinobacteria in NCBI.

### 2.2 Data loading and merging

The data is loaded into R and subsequently merged to a single object through the mmload function, which also extracts sequence length, GC content and tetranucleotide frequencies from the metagenome assembly. A standard workflow for data loading and merging is included as Load_data.Rmd.

### 2.3 Binning and validation

A number of functions are included in the mmgenome R package that facilitates effortless binning of metagenome contigs. The visualization functions wraps ggplot2 (Wickham, 2009) for easy manipulation and generation of publication quality plots. The main functions are described briefly below:

- *mmplot* Visualize contigs in 2D space using e.g. coverage and GC content and can integrate all associated data. For example, scale contigs by length, color by taxonomic classification and display paired-end read linkages (**Figure 1A**).

- *mmplot_network* facilitates visualization of paired-end linkages between contigs in a network graph. Any contig characteristic can be used for coloring (**Figure 1B**).

- *mmplot_locator* is used to interactively define a subspace in a *mmplot*. The coordinates are saved and used to extract the contigs within the subspace to a new mmgenome object using *mmextract.*

- *mmstats* calculates basic statistics on a mmgenome object. E.g. number of contigs, total length, N50, coverage, GC content and completeness estimates.

- *mmref* facilitates comparison of essential single copy genes to any group of genomes in NCBI in order to estimate completeness and contamination (**Figure 1C**).

- *mmexport* can be used to export the contigs of any mmgenome object as a fasta file.

### 2.4 Reassembly and finishing

Population genome bins can often be improved through reassembly. The script extract.fastq.for.reassembly.pl can be used to extract all paired-end reads associated with specific contigs.

To generate high quality population genome bins it is needed to manually inspect the binned contigs. The script circos.pl generates the raw data needed for circos plots, which integrates coverage, GC and paired-end linkage to assist in the final curation of the genome bin.

## 3 RESULTS

To demonstrate the capabilities of the mmgenome toolbox, we generated a time-series of four metagenomes from an enrichment laboratory scale reactor used to study enhanced biological phosphorus removal (Nielsen *et al.*, 2012). The dataset is included in the mmgenome R package, see supplementary information. An example workflow to extract an individual genome from the metagenome is shown in **Figure 1** and the basic statistics of the extracted population genome can be seen in **Table 1**. The complete workflow can be seen at http://madsalbertsen.github.io/mmgenome/ and recreated completely using the associated Rmarkdown file.

**Table 1.**
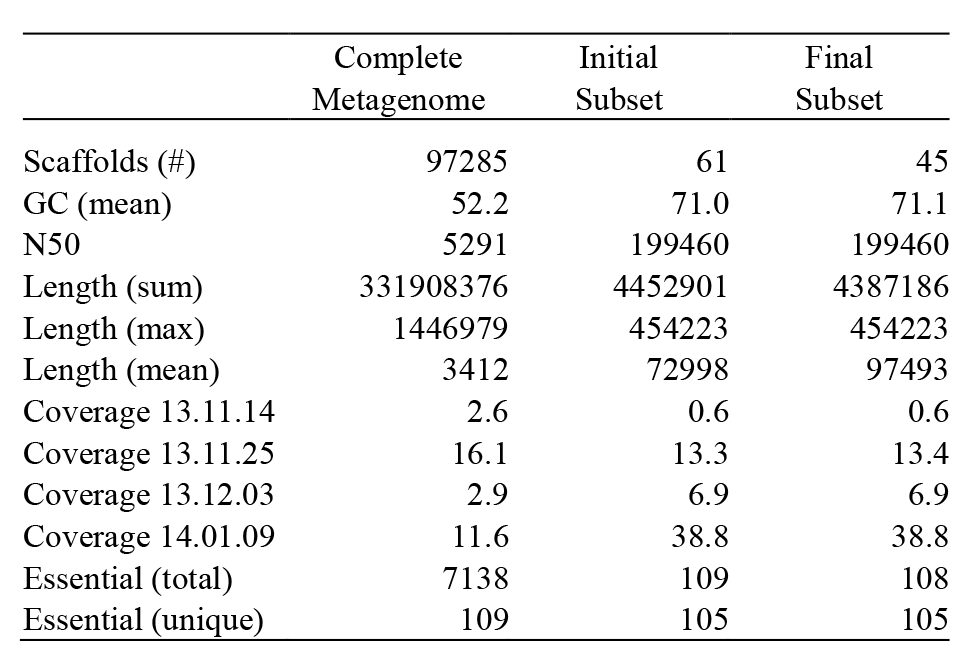
Statistics of the full metagenome and extracted genome bin calcu-lated using *mmstats*.

|  | Complete<br>Metagenome | Initial<br>Subset | Final<br>Subset |
| --- | --- | --- | --- |
| Scaffolds (#) | 97285 | 61 | 45 |
| GC (mean) | 52.2 | 71.0 | 71.1 |
| N50 | 5291 | 199460 | 199460 |
| Length (sum) | 331908376 | 4452901 | 4387186 |
| Length (max) | 1446979 | 454223 | 454223 |
| Length (mean) | 3412 | 72998 | 97493 |
| Coverage 13.11.14 | 2.6 | 0.6 | 0.6 |
| Coverage 13.11.25 | 16.1 | 13.3 | 13.4 |
| Coverage 13.12.03 | 2.9 | 6.9 | 6.9 |
| Coverage 14.01.09 | 11.6 | 38.8 | 38.8 |
| Essential (total) | 7138 | 109 | 108 |
| Essential (unique) | 109 | 105 | 105 |

## Acknowledgements

*Funding*: The project was funded by the Danish Strategic Research Council (Ecodesign-MBR) and the Villum Foundation.

